# Phenotypic architecture of sociality and its associated genetic polymorphisms in zebrafish

**DOI:** 10.1101/2021.07.29.454277

**Authors:** Claúdia Gonçalves, Kyriacos Kareklas, Magda C. Teles, Susana A. M. Varela, João Costa, Ricardo B. Leite, Tiago Paixão, Rui F. Oliveira

## Abstract

Sociality is often seen as a single phenotypic trait, but it relies on motivational and cognitive components implemented by specific causal mechanisms. Hence, these components may have evolved independently, or may have been linked by phenotypic correlations driven by a shared selective pressure for increased social competence. Furthermore, these components may be domain-specific or of general domain across social and non-social contexts. Here we have characterized the phenotypic architecture of sociality in zebrafish, which has been increasingly used as a model organism in social neuroscience. For this purpose, we have behaviorally phenotyped zebrafish from different wild type lines in four tests: social tendency, social and non-social recognition, and open-field test. Our results indicate that: (1) sociality has two main components that are independent from each other (social tendency and social recognition), hence not supporting the occurrence of a sociality syndrome; (2) both social traits are phenotypically linked to non-social traits (non-social exploration and non-social memory, respectively), forming two general behavioral modules, general inspection and general recognition, and suggesting that sociality traits have been co-opted from general-domain motivational and cognitive traits. Moreover, the study of the association between genetic polymorphisms (i.e. single nucleotide polymorphisms, SNPs) and each behavioral module further supports this view, since several SNPs from a list of candidate “social” genes, are statistically associated with the general inspection (motivational), but not with a general recognition (cognitive), behavioral module. The SNPs associated with general inspection are widespread across different chromosomes and include neurotransmitters, neuromodulators, and synaptic plasticity genes, suggesting that this behavioral module is regulated by multiple genes, each of them with small effects. Together, these results support the occurrence of general domain motivational and cognitive behavioral modules in zebrafish, which have been co-opted for the social domain.

**Author summary:** Social living has been considered one of the major transitions in evolution and it has been considered to act as a major selective force shaping the evolution of brain and behavior in animals. Sociality relies on two basic behavioral mechanisms: (1) the willingness to approach and be near others (aka social tendency); and (2) the ability to distinguish between others (aka social recognition) in order to adjust the behavior expressed during social interactions according to the identity of the interactant. There is an ongoing debate on to what extent these social abilities have specifically evolved in response to social living and are domain specific, or if they were selected as a broad response to cognitive demands and are of general domain. Here, we used zebrafish to test the domain-specific vs. general domain hypotheses and to assess the association of social tendency and social recognition with a set of candidate “social” genes (i.e. genes that have been linked to social behavior in other studies with different vertebrate species). We found that both social traits are not correlated to each other and are of general domain, and that only social tendency is associated with candidate “social” genes, suggesting that social tendency and social recognition are independent behavioral modules that rely on separate genetic architectures and that can evolve separately.

## Introduction

Sociality is ubiquitous among animals, with animal aggregations and the formation of social groups occurring across most animal taxa [1]. From a causal perspective sociality relies on two elementary behavioral mechanisms: (1) a motivation to approach conspecifics (social tendency) that leads to the formation of social groups; and (2) the cognitive ability to recognize different conspecifics (social recognition) that allows individuals to selectively adjust the expression of their behavior to different individuals they encounter. Given the fundamental role of these two behaviors for sociality, one can predict them to be selected together during social evolution, leading to a phenotypic correlation between them. On the other hand, each of these traits relies on different endophenotypes, with social tendency requiring a motivational (i.e. goal-directed) response, and social recognition requiring a cognitive ability (i.e. encoding, storing and recalling information about conspecifics to discriminate them), each implemented by different proximate mechanisms. For example, in mammals, social recognition is hippocampus-dependent (for a review see [2], whereas social tendency relies on mesolimbic reward circuits (e.g. [3–4]). Moreover, social recognition may reflect a general domain cognitive ability, that evolved to allow animals to discriminate different entities, social or not (e.g. edible vs. non-edible food), in the environment, rather than a domain-specific trait selected by sociality (e.g. [5–6]). In this case a phenotypic correlation would be expected between social recognition and non-social (e.g. object) recognition. Similarly, social tendency may reflect a general domain response to threat perception in the environment, since cohesiveness in animal aggregations is known to increase with perceived danger (i.e. aka defensive aggregation; e.g. rats: [7]; zebrafish: [8]). In this case a phenotypic correlation would be expected between social tendency and behavioral measures of anxiety/stress. Thus, the phenotypic architecture of sociality can be characterized by the pattern of phenotypic correlations among these behavioral traits.

The evolution of correlated traits can be explained by two alternative hypotheses: (1) the constraint hypothesis, that postulates the occurrence of shared proximal mechanisms such as a pleotropic effect of a gene, or a hormone with multiple target tissues; or (2) the adaptive hypothesis, that proposes that positive correlations between traits only occur in environments that favor them, such that selection can break apart maladaptive combinations of traits [9]. These two hypotheses generate different predictions that can be tested by comparing the patterns of correlated characters across different populations of the same species. The constraint hypothesis predicts traits to be correlated across populations irrespective of ecological conditions, whereas the adaptive hypothesis predicts correlations between traits to vary between populations depending on local conditions. Thus, these two scenarios also have different evolutionary consequences, with the correlated traits acting as evolutionary constraint in the first case and, the correlation being itself an adaptation in the latter. Although, this rational has been used to study the evolution of behavioral syndromes (aka personality) (e.g. [9]), to the best of our knowledge, it has not been applied yet to analyze the evolution of correlated social behavior traits. These two hypotheses can also be tested by assessing if the genetic architecture of correlated traits is shared or not. Given the complexity of social behavior traits, they are expected to be under the influence of multiple genes, with small effects of each of them. In fact, several genes involved in neurotransmission (e.g. dopamine, serotonin; [4;10–12]), neuromodulation (e.g. oxytocin; [13–15]) and synaptic plasticity mechanisms (e.g. neuroligins/ neurexins; [16–19]) have been reported to influence social behavior in multiple ecological domains across a wide range of vertebrate taxa. Moreover, these “social” genes are expressed in brain regions that together form an evolutionary conserved social decision-making network in vertebrates [20–21]. Therefore, the question is to what extent these candidate genes show specific or shared patterns of association with the motivational and cognitive components of sociality discussed above.

Enough variation in both social tendency and social recognition occurs across species and between individuals of the same species, which should allow to test the abovementioned hypotheses. The tendency to associate with conspecifics varies considerably among species, ranging from weakly social species, in which social interactions only occur at specific times (e.g. breeding), to highly social species, in which individuals stay all their lives in close proximity and interacting with others. Similarly, variation in social recognition ability also occurs across species, from basic levels of recognition (e.g. conspecific vs. heterospecific), to increasingly more elaborate ones with high degree of specificity (e.g. kin vs. non-kin; particular individuals) [22]. Moreover, variation in both social tendency and social recognition also occur within species, both intra- (e.g. with age and life-history stage) and inter-individually.

In this study we aim to characterize the phenotypic architecture of sociality in zebrafish (*Danio rerio*) by characterizing social tendency, social recognition and object recognition across multiple laboratory zebrafish populations that have evolved separately in captivity for multiple generations and by characterizing the genetic polymorphisms of candidate “social” genes associated with these behavioral traits. In zebrafish, isogenic lines are not viable due to inbreeding depression [23]. Hence, laboratory zebrafish populations differ from those of other model organisms in that they are recurrently outcrossed to maintain diversity [24]. As a result, laboratory zebrafish populations contain significant but varying levels of genetic diversity [25–26]). In parallel, zebrafish lines (e.g. AB, TU, WIK) have already been shown to vary in many behaviors, including locomotor activity, anxiety traits, stress reactivity, learning abilities and shoaling (e.g. [27–38]). The paralleled variation in genetic diversity [25–26] and several behavioral phenotypes, provides the rationale that constitutive genetic variation may contribute to the observed behavioral variability.

Here we specifically aim to test: (1) if there is an association between social tendency and social recognition, supporting the evolution of a sociality syndrome; (2) if social and non-social cognitive abilities (i.e. social vs. object recognition) are independent from each other, or if they co-vary supporting a general domain factor; (3) if there is an association between social tendency (or a putative sociality syndrome) and anxiety trait; (4) if the phenotypic correlations found are fixed or vary across lines (populations), in order to test the constraint vs. adaptive hypothesis; (5) to what extent the genetic architecture of each of these behavioral traits is shared or not, which would provide evidence for genetic pleiotropic effects underlying a putative sociality syndrome. For the latter, we have assessed the association between known single nucleotide polymorphisms (SNPs) in zebrafish for a set of candidate “social” genes (see methods for details) and each behavioral trait.

## Results

### Phenotypic architecture of sociality in zebrafish

Scores of social tendency (i.e. preference for shoal over empty tank), as well as object and conspecific discrimination scores (i.e. preference between a novel and a familiar stimulus) were all significantly different than chance for individuals of both sexes and for all lines tested (one-sample *t*-test: *μ* ≠ 0.5, *P* < 0.001; see Table S1; Fig 1f-h), indicating that social affiliation and social and object recognition abilities are present in males and females across zebrafish lines.

**Fig 1.**
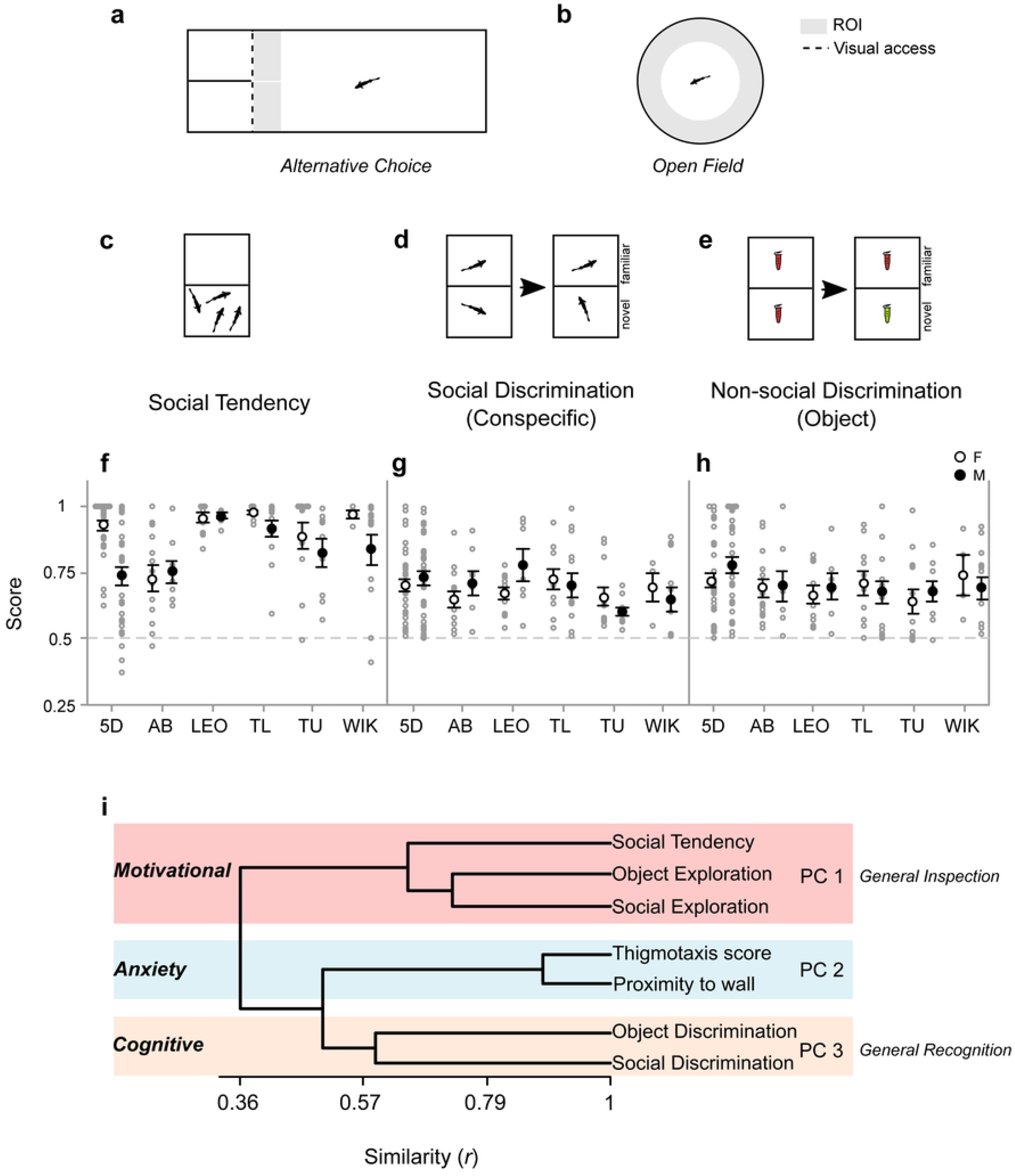
Social and associated behaviors in zebrafish. (**a**) Across lines, a two alternative-choice set-up was used to measure social preference and recognition abilities (**b**) and an open field test of measuring anxiety-driven thigmotaxis towards the periphery and edge-orienting. Regions of interest (ROI) were set within 1 standard body-length from target locations or stimuli. (**c**) Social tendency was measured by interaction preferences towards a shoal. Recognition in both the social (**d**) and non-social (**e**) context were measured by the ability to discriminate between a familiar and a novel stimulus. (**f**) Males (full circles) and females (open circles) of all lines (5D, AB, LEO, TL, TU, Wik) exhibited above chance (dashed line) preference for shoal over an empty tank (social tendency, **f**) and discrimination between a novel and familiar stimulus in both a social (conspecific; **g**) and non-social (object; **h**) context [bars indicate 95% CI]. Behavioral measures exhibited different degrees of correlation (*r*), illustrated in the cladogram as degrees of association (**i**), based on which factor analysis revealed three principal components (PC): PC1 aggregates social tendency and social and object exploration corresponding to a motivational component of sociality; PC2 aggregates thigmotaxis and (i.e. proportion time in periphery) and edge-orienting (distance to wall) measured in the open field test, corresponding to an anxiety component; PC3 aggregates object and social discrimination, corresponding to a general-domain cognitive component.

To assess the phenotypic architecture of sociality we performed factor analysis using principal component extraction followed by varimax rotation, based on the correlation matrix between measures extracted from the 4 separate tests of social and associated behaviors (sampling adequacy: KMO > 0.5; sphericity: Bartlett’s*χ^2^_21_* = 253.76, *P* < 0.001; determinacy of multicollinearity: *ρ* = 0.754). The analysis identified 3 components (C) with eigenvalues ≥ 1 (Fig 1i and Table 1). C1 shows a strong loading of social tendency measured in the social preference test and of social and object exploration measured in the social and object discrimination tests, respectively, suggesting the occurrence of a general inspection behavioral module that is expressed both in social and non-social contexts. C2 shows a strong loading of thigmotaxis and edge-orienting measured in the open-field test, corresponding to an anxiety behavioral module. Finally, C3 shows a strong loading of object and social discrimination, measured in the object and social discrimination tests, respectively, suggesting the occurrence of a general recognition behavioral module that is expressed both in social and non-social contexts.

**Table 1.** Loadings extracted by the varimax rotaton of principal components from the correlation matrix of behaviors across tests, for zebrafish of all lines. Bold type indicates the strongest contributors (coefficient > 0.5) to each component (C).

| Test | Behavior | Parameters | Component loadings <sup>a</sup> |  |  | Comm. <sup>b</sup> |
| --- | --- | --- | --- | --- | --- | --- |
|  |  |  | C 1<br>*General Inspection* | C 2<br>*Anxiety* | C 3<br>*General Recognition* |  |
| SP | Social tendency | Proportion total ROI time spent with shoal | **0.725** | -0.033 | 0.180 | 0.559 |
| SR | Social discrimination | Proportion total ROI time spent with preferred conspecific | 0.113 | -0.031 | **0.852** | 0.739 |
|  | Social Exploration | Proportion test time spent in ROI of both conspecifics | **0.762** | 0.085 | -0.224 | 0.639 |
| OR | Non-social discrimination | Proportion total ROI time spent with preferred object | -0.363 | 0.139 | **0.619** | 0.535 |
|  | Non-social exploration | Proportion test time spent in ROI of both objects | **0.754** | -0.042 | -0.111 | 0.583 |
| OF | Thigmotaxis | Proportion time spent within ROI of periphery, after first entry (control for initial freezing in center) | 0.107 | **0.932** | 0.084 | 0.888 |
|  | Edge-orienting | Average distance from wall (cm) | 0.112 | **-0.941** | 0.009 | 0.897 |
| *Eigenvalue <sup>c</sup>* |  |  | *1.845* | *1.784* | *1.211* | *4.840* |
| *% Variance explained* |  |  | *0.264* | *0.255* | *0.173* | *0.691* |

<sup>a</sup> Correlation between components and variable values
<sup>b</sup> Communalities: Proportion of variable variance explained by all components
<sup>c</sup> Variance of transformed data used for each component

To test if the behavioral modules described above (i.e. General inspection, General recognition, and Anxiety) can evolve differently from each other in each zebrafish line – which represent different laboratory populations established by different wild type founders and that have evolvedindependently from each other in somewhat similar lab conditions – we computed correlation matrices between individual scores for each module (varimax rotated PC scores)for each of the different zebrafish lines. We then used the quadratic assignment procedure (QAP) correlation test to compare the correlation matrices of the different lines. The results identified a single significant negative correlation (r = −0.9988, p= 0.0002) between 5D and WIK correlation matrices. Thus, none of the correlation matrices were similar between each other (Fig 2), supporting the adaptive hypothesis, that predicts different patterns of phenotypic correlations in different populations.

**Fig 2.**
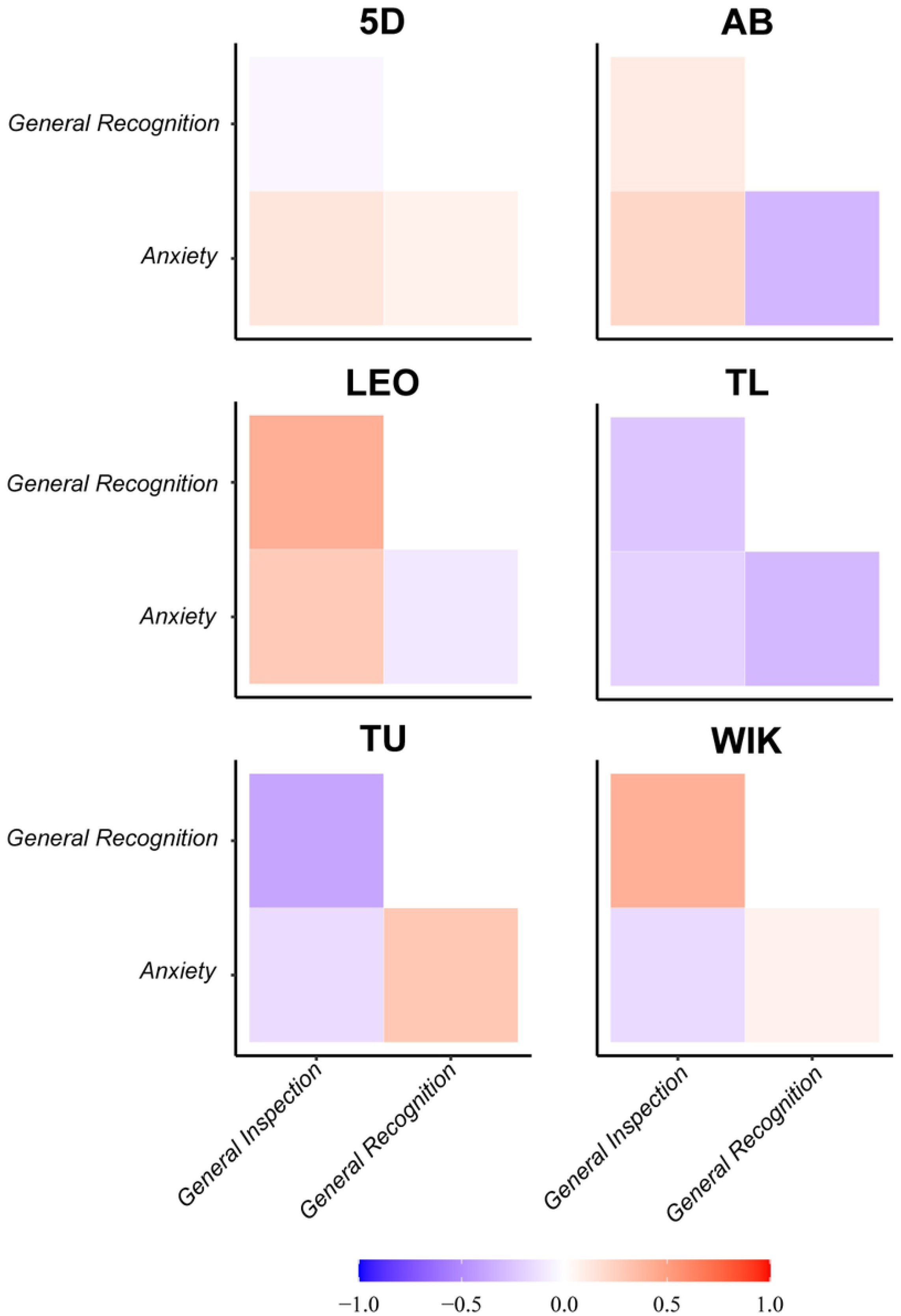
Phenotypic correlation matrices. Phenotypic correlation matrices for the behavioral components of General Inspection, General Recognition and Anxiety across six different zebrafish laboratory lines (i.e. populations). Color code represents correlation (*r*) values.

### Genetic polymorphisms associated with behavioral modules

To assess if the different behavioral modules identified above were linked by a shared genetic architecture, we have investigated the association between a set of genetic polymorphisms (SNPs) in a list of candidate “social” genes and each of the measured behaviors and PC behavioral modules. Given the fact that we have phenotyped individuals from 6 different wild type lines, we checked for structured genetic variation by computing the genetic distance between the phenotyped individuals for the SNPs under study. We found that genetic variation for the SNPs of interest is highly structured with individuals from the same wild type lines clustering together (Fig 3a). Therefore, we have used the line as a covariate in the model that assessed the association between each SNP an each of the behavioral modules.

**Fig 3.**
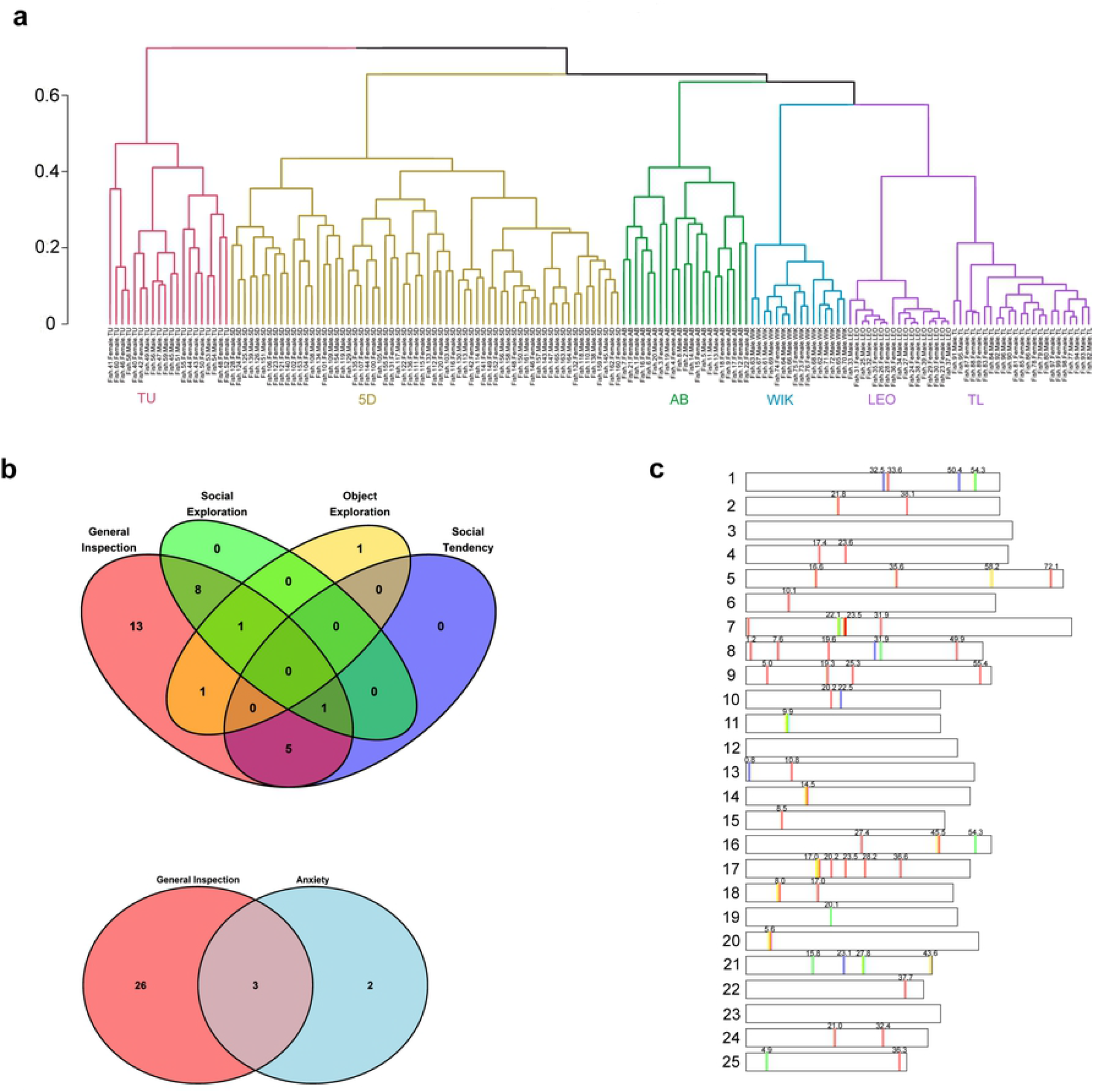
Genetic clustering and behavioral associations. (**a**) Hierarchical clustering of genetic distances (Jaccard distance) between the sampled individuals indicates the occurrence of 5 major clusters that overall match the 6 wild type lines used (pink cluster: TU; gold cluster: 5D; green cluster: AB; blue cluster: Wik), with LEO and TL included in the purple cluster but subsequently segregated from each other in two lower order clusters. (**b**) Venn diagrams representing the number of SNPs the General Inspection component shares with its constitutive behaviors (social tendency, social exploration, and object exploration) and the Anxiety component. (**c**) Chromosome mapping of the SNPs that are significantly associated with the General Inspection component and its constitutive behaviors, following the color code used by the Venn diagrams, and with the position of each SNP on each chromosome is given in bp.

Out of the 132 SNPs that showed variation in our sampled individuals, 52 (which mapped to 29 genes) were significantly associated with General Inspection, none with General Recognition and 8 (which mapped to 5 genes) with Anxiety (Table 2). Regarding the 3 behaviors that loaded to the General Inspection behavioral module, 6 SNPs (mapping to 6 genes) were associated with social tendency, 11 (mapping to 10 genes) with social exploration, and 3 (mapping to 3 genes) with object exploration. Of these 20 SNPs associated with these behaviors that load to General Inspection, only one (mapping to the serotonin receptor gene 5HTR 2cl2) is not also associated with General Inspection (Fig 3b; Table 2). Moreover, of the 29 SNPs associated with General inspection, 16 are also associated at least with one of the behaviors that constitutes these behavioral module (Fig. 3b; Table 2). However, there is a reduced overlap between the SNPs associated with these different behaviors: only one SNP affects both social tendency and social exploration (matching the gene 5HTR-1aa), and only another SNP affects both social exploration and object exploration (matching the gene 5HTR-2cl1) (Fig. 3b; Table 2).

**Table 2.**
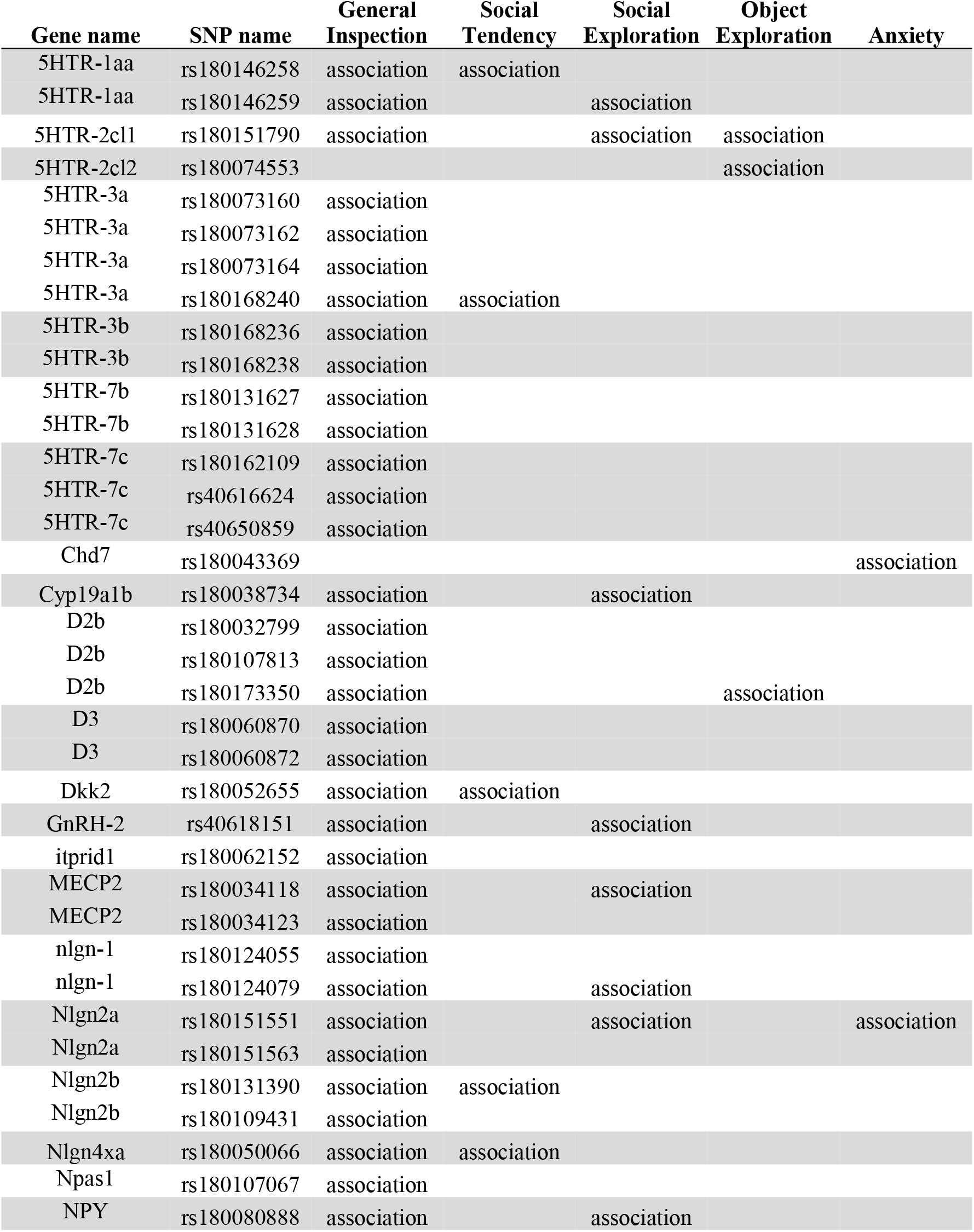
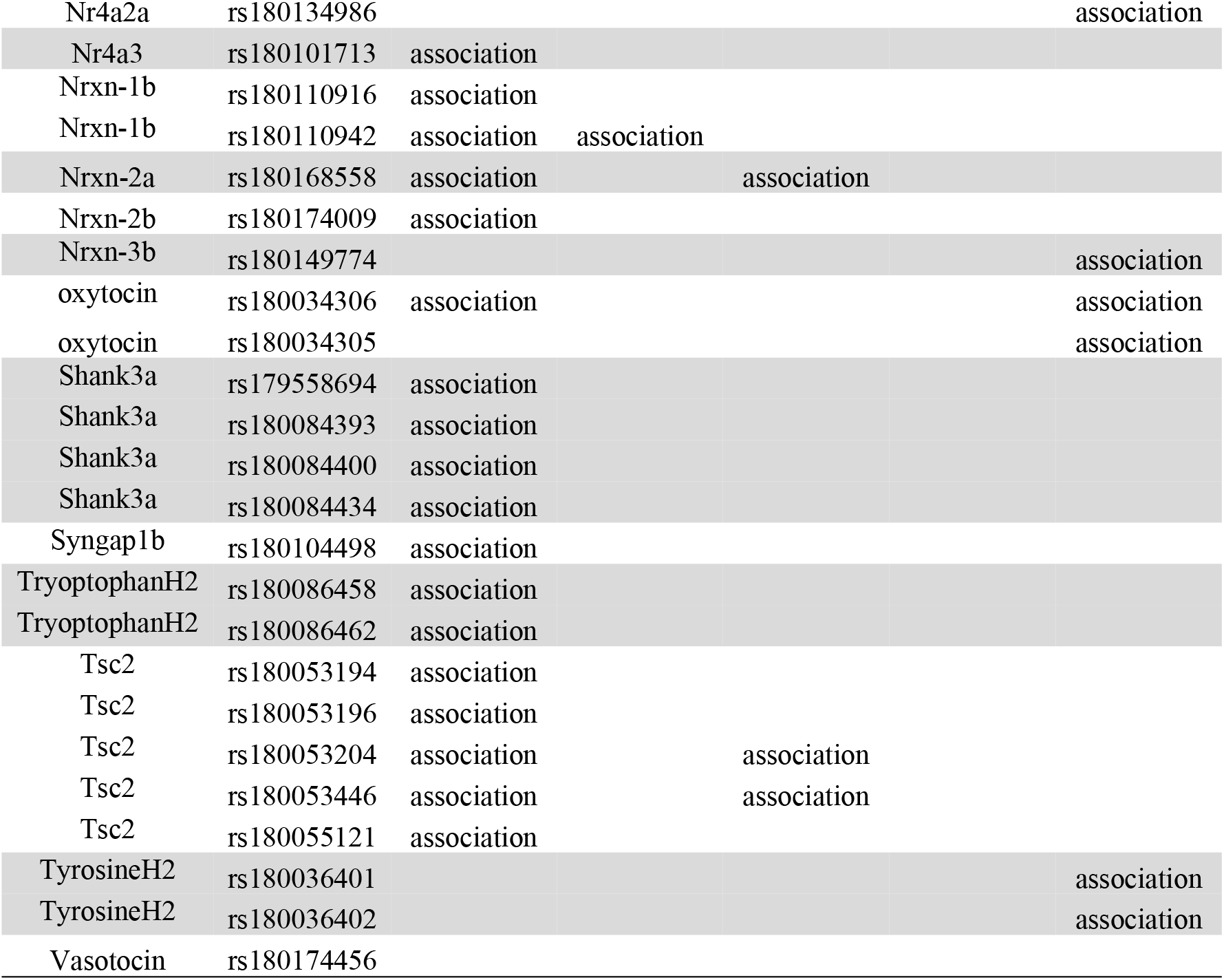
Lists of genes with SNPs associated with the behavioral modules General Inspection (and its contributing behaviors) and anxiety. Association = the SNP is associated with that behavior. Abbreviations: 5HTR=serotonin receptor; D = dopamine receptor; Cyp19a1b = cytochrome P450, family 19, subfamily A, polypeptide 1b; Nrxn = neurexin; Nlgn = neuroligin; Npas1 = Neuronal PAS Domain Protein 1; NPY = neuropeptide Y; Nr4a3= nuclear receptor subfamily 4, group A, member 3; Nr4a2a= nuclear receptor subfamily 4, group A, member 2a; Dkk2 = dickkopf WNT signaling pathway inhibitor 2; itprid1 = ITPR interacting domain containing 1; MECP2 = methyl CpG binding protein 2; Syngap1b = synaptic Ras GTPase activating protein 1b; Tsc2 = TSC complex subunit 2; Chd7 = chromodomain helicase DNA binding protein 7.

The SNPs associated with the General Inspection behavioral module are widely distributed across the zebrafish genome being absent only from chromosomes 11, 12, 19, 21 and 23 (Fig 3c). However, one can find SNPs associated with behaviors that load to General Inspection module in some of these chromosomes; SNPs associated with social exploration in chromosome 11, 19 and 21; and SNPs associated with social tendency and with object exploration in chromosome 21 (Fig 3c).

The list of SNPs associated with the General Inspection module include genes involved in neurotransmission (e.g. serotonin and dopamine receptors), neuromodulation (e.g. NPY, oxytocin), synaptic plasticity (e.g. neuroligins, neurexins) and epigenetic marking (e.g. methyl CpG binding protein 2) (see Fig 4 for arbitrarily selected illustrative examples).

**Fig 4.**
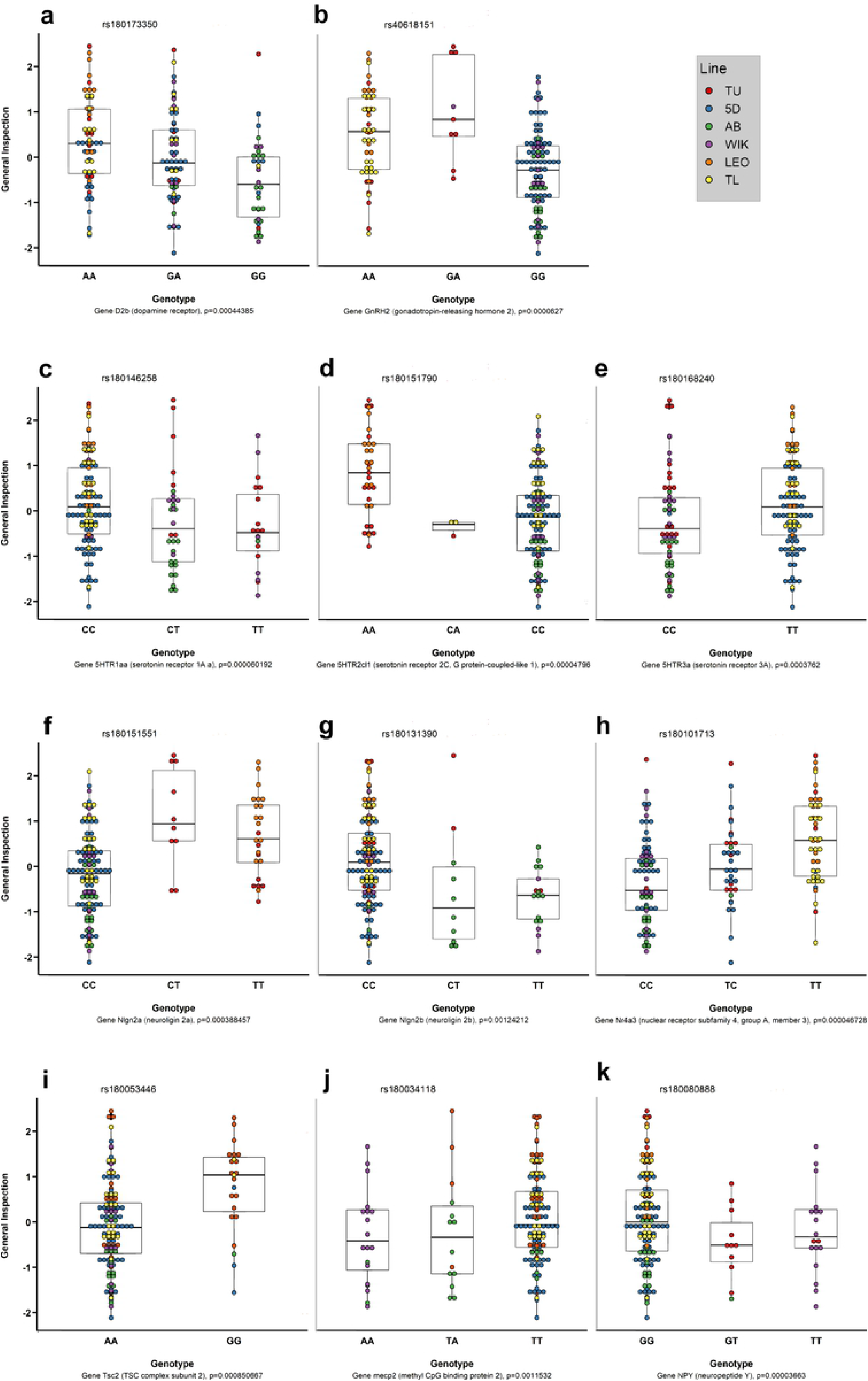
Illustrative examples of SNPs associated with the General Inspection component: (**a**) D2b; (**b**) GnRH2; (**c**) 5HTR1aa; (**d**) 5HTR2d1; (**e**) 5HTR3a; (**f**) Nlgn2a; (**g**) Nlgn2b; (**h**) Nr4a3; (**i**) Tsc2; (**j**) MECP2; (**i**) NPY. Individuals of the different lines are represented by different colors according to color code indicated in the figure legend.

### Sex and line differences in behavioral modules

Although it was not the central question of this study, the occurrence of sex and line differences in the expression of the behavioral modules identified above can be informative when choosing lines to run specific behavioral tests in zebrafish. Therefore, we have also tested for both sex and line differences in General Inspection, General Recognition and Anxiety (Fig 5).

**Fig 5.**
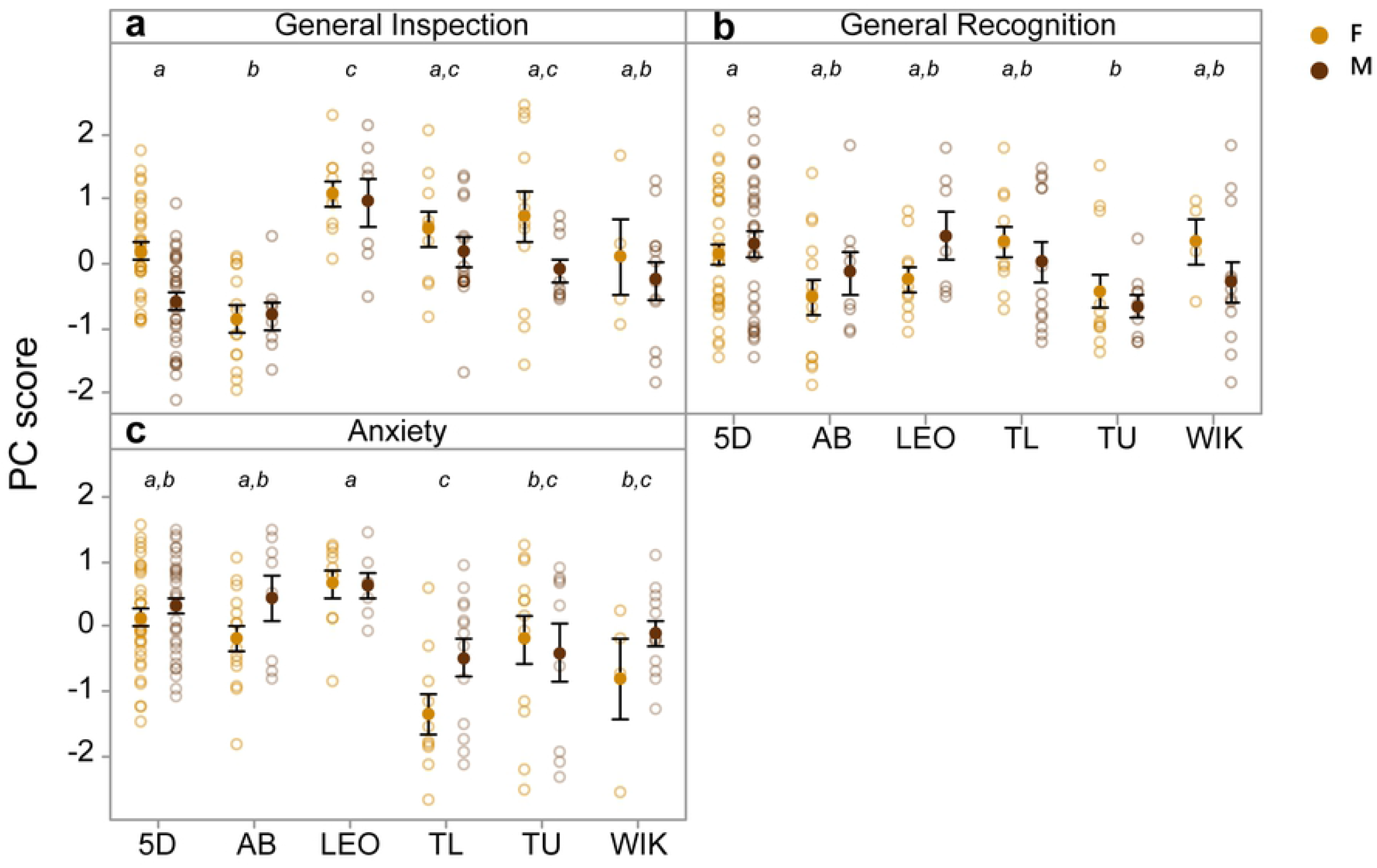
Comparison of General inspection, General Recognition and Anxiety across lines and sexes. Orange=females; black = males. Individual scores ranged between −2 and 2. Bars indicate 95% Confidence Intervals with Bonferroni correction and means that do not share a letter are significantly different [*P* < 0.05].

General Inspection presented a main effect for both sex and line, but not for the interaction between them (sex: *F* _1, 163_ = 6.70, p < 0.05; line: *F* _5, 163_ = 11.04, p < 0.001; sex*line: *F* _5, 163_ = 1.22, p = 0.304), with females having significantly higher scores than males, and LEO having the best performance which was significantly higher than that of Wik, 5D, and AB, with 5D being also significantly higher than AB.

General recognition only presented a significant main effect for line (sex: *F* _1, 163_ = 0.01, p = 0.994; line: *F* _5, 163_ = 2.64, p < 0.05; sex*line: *F* _5, 163_ = 1.04, p = 0.396), with 5D having significantly higher performance than TU, and all other lines having intermediate, and not statistically different, performances.

Anxiety presented a main effect for both sex and line, but not for the interaction between them (sex: *F* _1, 163_ = 4.75, p < 0.05; line: *F* _5, 163_ = 8.32, p < 0.001; sex*line: *F* _5, 163_ = 1.21, p = 0.396), with males having significantly higher scores than females, and among lines LEO having the highest score which was significantly higher than that of TU, Wik, and TL, with the latter (TL) being significantly lower than 5D and AB.

## Discussion

In this study we have characterized the phenotypic architecture of sociality in zebrafish. We have behaviorally phenotyped males and females of six different wild type laboratory lines in 4 behavioral tests (social tendency, social and object discrimination and open-field) and showed that social tendency (i.e. preference to associate with conspecifics) and the ability to discriminate between conspecifics (social recognition) is present in both sexes of all lines tested. A factor analysis identified three main behavioral modules: (1) general inspection, which includes social tendency measured in the social preference test and social and object exploration, measured in the social and object discrimination tests, respectively; (2) general recognition, which includes social and object discrimination, measured in the social and object discrimination tests, respectively; and (3) anxiety, which include the behavioral measures of thigmotaxis and edge-orienting taken in the open field test. Therefore, the motivational (social tendency) and cognitive (social recognition) aspects of sociality are not phenotypically correlated, a result that does not support the occurrence of a sociality syndrome, which could be predicted by shared selective pressures on these two traits for the evolution of sociality. Moreover, the fact that both social tendency and social recognition are phenotypically correlated with similar non-social behaviors (i.e. object exploration and object recognition, respectively), integrating two general-domain behavioral modules (general inspection and general recognition), supports the hypothesis that these behaviors are not domain specific and have been evolutionarily co-opted from general-domain motivational and cognitive traits. These results agree with a recent study showing that in zebrafish both social recognition and object recognition, but not social tendency, are oxytocin-dependent, hence suggesting a common proximate mechanism indicative of a general-domain cognitive trait [39]. Finally, it is worth noting that even though sociality has been proposed to be promoted by predator pressure as a defensive mechanism [40], anxiety forms an independent behavioral module from those where social traits are included.

Even with the motivational and the cognitive components of sociality being part of two different behavioral modules, a shared selective pressure on both for the enhancement of social competence could result in a physiological linkage between the two behavioral modules; for example, due to the evolution of a common neuromodulator that phenotypically integrates the independent neural mechanisms underlying general inspection and general recognition. In fact, even though that social affiliation and social memory have been shown to rely on separate neural circuitry, some neuromodulators, such as oxytocin have been shown to regulate both mechanisms (e.g. [41–42]), opening the possibility for the evolution of physiological constraints that phenotypically link these two domains. We tested the constraint hypothesis, which predicts traits to be correlated across populations irrespective of ecological conditions [9;43], in our data set by comparing the matrices of phenotypic correlations among the three behavioral modules extracted from the factor analysis across the six wild type lines used in this study. Given that these wild type laboratory lines have been established independently from different founders collected in the wild and have been evolving independently from each other in similar stochastic lab environments, they can be seen as independent representative populations of this species (despite living in artificial environments). Contrary to the prediction of the constraint hypothesis, the phenotypic correlation matrices were not similar between any pair of zebrafish laboratory lines studied. In fact, there was only one significant QAP correlation between the 5D and Wik matrices, but it was a negative correlation suggesting an asymmetric structure of the matrices. Therefore, our data supports the alternative adaptive hypothesis, that proposes that positive correlations between traits are the result of historical selection favoring particular trait combinations (i.e. selection-induced linkage disequilibrium [44–45], such that the evolution of different combinations between the different behavioral modules is not physiologically or genetically linked, and hence divergence of the correlation matrices between populations is unconstrained.

The study of the association between a set of genetic polymorphisms (SNPs), in candidate genes that have been implicated in social behavior in vertebrates (“social genes”), and the behavioral modules that emerged from our factor analysis indicates that only the general inspection (motivational) module is associated with SNPs in the “social genes”, further supporting the lack of genetic linkage between this module and the general recognition (cognitive) module. Thus, the “social genes” studied here seem to be associated with a general domain motivational component of social behavior, rather than with a general domain cognitive component, which probably relies on memory related genes not included in our “social genes” list. Moreover, our results also indicate a low overlap in the genetic polymorphisms association (3 out of 29 SNPs) between the general inspection and the anxiety modules, which suggests that despite these two behavioral modules relying on motivational mechanisms they have significantly different genetic architectures.

Interestingly, all except one of the genetic polymorphisms (5HTR 2cl2) associated with the 3 behaviors that loaded to the General Inspection behavioral module, are also associated with this behavioral module indicating an agreement between phenotypic (i.e. behavioral correlations) and the genetic (i.e. genetic polymorphisms) data supporting the occurrence of this behavioral module. The genetic polymorphisms associated with these behaviors include neurotransmitter and neuromodulator systems known to modulate motivational states, such as serotonergic (social tendency is associated with, 5HTR1aa, 5HTR3a and social exploration with 5HTR-1aa, 5HTR-2cl1) and neuropeptidergic pathways (social exploration is associated with GnRH2 and NPY), as well as genes involved in synaptic plasticity, such as the neuroligin/neurexin system (social tendency is associated with Nrxn1b, Nlgn2b, and Nlgn4xa, and social exploration with Nlgn1, Nlgn2a, and Nrxn2a) and epigenetic marking (social exploration is associated with the methyl CpG binding protein 2). On the other hand, the genetic polymorphisms associated with object exploration include less “social genes” (only 3), which are restricted to the serotonergic and dopaminergic neurotransmitter pathways (5HTR-2cl1, 5HTR-2cl2, D2b). Thus, even within a behavioral module it is possible to observe a significant partitioning of the genetic associations with the different component traits of that module. This conclusion is further supported by the fact that there are only 2 SNPs, in the same gene (5HTR-1aa), that are associated both with social tendency and social exploration, and only another SNP in one gene (5HTR-2cl1) associated both with social and object exploration.

The SNPs associated with the General Inspection behavioral module are distributed across 20 of the 25 chromosomes that constitute the zebrafish genome, being absent only from chromosomes 11, 12, 19, 21 and 23. However, one can find SNPs associated with behaviors that load to the general inspection module in chromosomes that do not contain SNPs associated with the behavioral module itself (e.g. SNPs associated with social exploration in chromosome 11,19 and 21, and the SNPs associated with social tendency and with object exploration in chromosome 21). In a previous study that aimed to identify quantitative trait loci (QTL) in zebrafish for behavioral and morphological traits, QTLs for social tendency have been identified when using one of the two statistical methods used (genetic algorithm mapping vs. interval mapping) in chromosomes 18 and 24 [46]. In our study, variation in social tendency is associated with SNPs located in chromosomes 1(#2), 8, 10, 13 and 21. However, the General Inspection module, where social tendency is included, has associated SNPs on chromosomes 18 and 24. Hence, this mismatch between the QTL results and our results presented here can be due either to a false detection of these QTLs by the genetic algorithm mapping method, given the lack of support from the interval mapping method in the previous study, which led the authors not to claim these QTLs themselves [46]; or to an indirect association through the link between social tendency and the general inspection module. Either way, our results show that the SNPs associated with both the general inspection module and the behaviors that constitute this module are widespread across the genome, supporting a many gene (each with small effects) genetic architecture for these traits.

## Methods

### Ethics statement

All experimental procedures were reviewed by the institutional internal Ethics Committee at the Gulbenkian Institute of Science and approved by the National Veterinary Authority (Direção Geral de Alimentação e Veterinária, Portugal; permit number 0421/000/000/2017).

### Zebrafish lines and housing conditions

Zebrafish were raised in the Fish Facility of the Gulbenkian Institute of Science under laboratory conditions. The following lines were used in this study:

1. AB, was established by George Streisinger and Charline Walker in the Oregon labs, from two lines, A and B, purchased by George Streisinger at different times from a pet shop in Albany, Oregon, in the late 1970s. The original A and B lines probably originated from a hatchery in Florida. The AB line has been screened for lethal-free embryos by in vitro fertilization and selected females subsequently used to establish the current AB line [47–48]. This procedure reduced the number of lethal mutations in this line, which has been used as the primary background for most of the transgenic and mutant lines that are currently available.
2. TU (Tuebingen), originated from a pet store in Tuebingen and was selected during the 1990s at the Max-Planck in Tuebingen to remove embryonic lethal mutations from the background before being used by Sanger for the zebrafish sequencing project [49–50].
3. WIK (Wild India Kolkata), was derived from a wild catch of a single pair in India, near Kolkata. The WIK line is very polymorphic relative to the TU line and was first described as WIK11 [51].
4. TL (Tüpfel long fin), was derived from a cross between an AB with a spotted phenotype and a TU resulting in a long-finned phenotype. This line is homozygous for leot1, a recessive mutation causing spotting in adult fish (aka tup), and for lofdt2, a dominant mutation causing long fins.
5. 5D (5D Tropical), was derived at Sinnhuber Aquatic Research Laboratory (SARL) at Oregon State University in 2007, from a commercial breeding facility (5D Tropical Inc., Florida), to generate a *Pseudoloma neurophilia* (Microsporidia) free line [52].
6. LEO (Leopard), is a wild type line commonly available in pet shops, which displays a spotted adult pigment pattern instead of striped. This line is homozygous for a spontaneous mutation in the gene leopard (leo), leot1 [53–55].

A total of 164 experimentally naive adult zebrafish of both sexes, aged 6-8 months, were used in this study as focal subjects (AB: M = 8, F = 14; TU: M = 9, F = 12; WIK: M = 12, F = 4; TL: M = 13, F = 10; LEO: M = 7, F = 10; 5D M = 32, F = 33). Focal fish were raised and housed separately from fish used as stimuli to prevent effects of prior familiarity. Fish used as stimuli were of the same line as the focal fish. Housing was in groups of 35 fish kept in 3.5 L aquaria of a recirculating system (ZebraTec, 93 Tecniplast), with water parameters set at 27-28 °C, 7.5 ± 0.2 pH, ~ 900 μSm, and <0.2 ppm nitrites, <50 ppm nitrates and 0.01-0.1 ppm ammonia. Daily photoperiods were alternated between 14h light and 10h dark and feeding occurred twice-daily and included a combination of live (*Paramecium caudatum; Artemia salina*) and processed dry food (GEMMA Micro).

### Experimental setup and procedures

The behavior of each experimental fish was assessed in four different tests: (1) a shoal preference test to measure social tendency; two one-trial recognition tests using either objects (2) or conspecifics (3) as stimuli to measure non-social and social recognition/exploration, respectively; and (4) an open-field test to measure the anxiety trait. Excluding the open-field, all setups included an experimental tank (30 L x 15 W x 15 H cm) and two adjacent tanks (15 L x 15 W x 7.5 H cm) with a stimulus-holding compartment having a viewing side of 10 cm in the shoal preference test and of 5 cm in the social and object recognition tests, the difference accounting for the different visual target areas offered by a shoal *vs*. an individual or an object. Water depth was kept constant at 9cm in all tanks (Fig 1a). For the open-field test, a round tank with a 22 cm diameter was used and water level was kept at 6 cm depth (Fig 1b).

All tests occurred during the light period between 09:00 and 19:00, before which fish were kept overnight in an aquarium with individual compartments for identification purposes. These compartments were separated by fine mesh that allowed visual and chemical access to neighbors and minimized stress from isolation. In the experimental tanks, external stimuli were visually blocked by opaque, non-reflective stickers and opaque covers obscured adjacent stimulus containers prior to the onset of recordings during the shoal preference and recognition tests. Behavior during tests was recorded using black and white mini surveillance cameras (Henelec 300B) suspended above the experimental tank and relaying the image to a laptop kept at a distance to reduce disturbance of fish by the experimenter. During recording, lighting in the room was kept at conditions that reduce water-surface reflection in the videos, and extra lighting was provided by an infrared lightbox placed under the experimental tank in order to facilitate video tracking during the data collection stage. Between tests, water in the experimental tank was changed to eliminate olfactory cues.

Before tests, focal fish were netted from their individual overnight compartment and immediately placed in the experimental tank. For the shoal preference test, fish were first given 10 min to acclimatize to the experimental tank and then tests were initiated by removing the opaque covers and allowing fish visual access to the two adjacent containers, one empty (control) and the other holding a mixed-sex shoal (Fig 1c), for 10 min. The side of presentation of each stimulus was counterbalanced between focal individuals to control for side biases. Recognition tests were comprised of two phases: an acquisition phase and a probe-test phase, and the experiment included a 10 min initial acclimation period before the acquisition phase and a 10 min interval before the probe-test phase. Both phases were initiated by removing opaque covers and allowing fish visual access to two adjacent containers. During the acquisition phase, animals were presented with two novel stimuli for 10 min: two conspecifics for the social test and two objects (0.5 ml eppendorf tubes of the same color) for the non-social test. During the probe test, animals were presented with one of the stimuli from the acquisition phase (familiar) and a novel stimulus (a new conspecific or a differently colored eppendorf tube) for 10 min (Fig 1d). For the non-social recognition test, the size of the eppendorf tubes was matched to the average zebrafish size to control for size-dependent prey or predator directed responses and, based on preliminary preference tests, were colored with colors of equal preference by the fish (either green or red for all lines, except for LEO that instead show no preference between purple and blue). The side of each stimulus (novel or familiar) during probe tests was counterbalanced across animals, to control for side biases, and the color used for the familiar or novel stimulus was randomized, to control for color biases (Fig 1e). For the open-field test (Fig 1b), animals were placed in the center of the circular tank and recorded for 10 min.

Videos were analyzed using a commercial video-tracking software (EthoVision XT, Version 11.5, Noldus Information Technology) and behavioral measures were extracted from each test. Regions of Interest (ROI) marked were kept at an average body length distance from the target location (grey regions in Fig 1a-b). Social tendency during the shoal preference test was quantified by the proportion of time in ROIs spent near the shoal, social and non-social discrimination during the conspecific and object recognition tests was measured by the proportion of time in ROIs spent near the preferred stimulus (familiar or novel), while the overall time spent in ROIs near both stimuli was used as a measure of exploration. Anxiety in the open field test is typically exhibited by thigmotaxis (i.e. the propensity to avoid exposed areas), which was measured as the proportion of time spent within the ROI near the periphery following first entry (to control for any initial freezing in the center), while the average distance (in cm) from the wall was used to quantify the edge or wall orienting tendency associated with fear-induced thigmotaxis [56].

### Genetic polymorphisms analysis

At the end of the behavioral phenotyping, animals were anesthetized by immersion into an ethyl 3-aminobenzoate methanesulfonate salt solution (MS222) 100-200mg/L, a fin clip collected from the caudal fin of each experimental fish, and preserved in a digestion mix (PK, 10 mg/ml, Lysis solution [Fermentas #K0512], TE buffer) until further processing. Subsequently, DNA was extracted from preserved fin clips using DNA Extraction kit (Fermentas #K0512) with some adjustments to the protocol provided by the manufacturer. Briefly, samples were thawed at room temperature and placed in a thermomixer for approximately 20h with shaking (700 rpm) at 50°C. After, chloroform was added in a 1:1 ratio and the samples gently mixed by inversion. Samples were then centrifuged at 18506 g (13200 rpm in Eppendorf 5430R centrifuge) for 7 min and the upper aqueous phase transfer to a new 1.5 ml tube. 800 μl (720μl H_2_O + 80μl of precipitation solution [Fermentas #K0512]) was added to each tube, mixed gently by inversion for 2 min and centrifuged again for 10 min at 18506 g. The supernatant was removed, the DNA pellet dissolved in 100μl NaCl 1.2M solution [Fermentas #K0512], and 300μl of freezer cold 100% ethanol (−20°C) was added to allow DNA to precipitate over night at −20°C. In the day after, samples were centrifuged for 10 min (18506 g) and the ethanol removed. To wash the pellet, 200 μl of freezer cold 70% ethanol was added to each sample and centrifuged for 10 min (18506 g). Finally, the pellet was allowed to dry for 15-30min at 37°C and 30μl of DNAse-free sterile H2O was added. To access the concentration and quality of the DNA, samples were quantified in the Nanodrop (Thermo Scientific, Nanodrop 2000) and the ratios 260/280 and 260/230 listed.

We built a list of candidate genes to test their association with the behavior traits, based on evidence from the literature for their involvement in the regulation of social behavior. This gene list included genes for: neurotransmitter systems (e.g. dopamine, serotonin), neuromodulators (e.g. oxytocin, AVT, NPY), neuroplasticity (e.g. bdnf, neurexins, neuroligins), and genes linked to autism (e.g. shank3a). To obtain candidate SNPs for the genes of interest, all germline variations from this species were downloaded from http://ftp.ensembl.org/pub/release-104/variation/gvf/danio_rerio/ in the form of a GVF file. The GVF file was filtered to keep only SNPs in locus of interest and which evidence was sustained by frequency observations to increase probability of variation. Sequences were extracted with Ensembl’s Biomart tool using the “Zebrafish Short Variants (SNPs and indels excluding flagged variants) (GRCz11)” dataset. Several iterations of Assay Design 4.0 (Agena Biosciences), which designs multiplexed MassEXTEND® assays for Mass Spectrometry detection, were run to accomplish an even distribution on the genes of interest. Four multiplexes were designed with 38, 36, 35 and 35 assays. Agena Biosciences iPlex(®) Kit, MassARRAY(®) platform and Typer software v.4 were used following manufacturer’s standard protocols and procedures, for the genotyping reactions, acquisition of genotypes and inspection of results, respectively. 139 SNPs in locus of interest were successfully sequenced, but we had to remove 7 for lack of variation between the 164 tested zebrafish (the final list of SNPs is available in Table 2).

### Statistical Analysis

In order to confirm that all lines express social tendency and are able of social and object recognition, one-sample *t*-tests (*μ* ≠ 0.5 vs. >0.5) were used to test if the scores of social tendency, object discrimination and social discrimination were significantly different from chance levels for each sex and for each line. Next, we extracted behavioral modules that aggregate correlated behaviors by carrying out a factor analysis using principal component extraction (PCA) followed by varimax rotation, based on the correlation matrix of all behavioral measures (social tendency, social discrimination, social exploration, object discrimination, object exploration, thigmotaxis and edge-orienting). The analysis identified three main components (Cs) to which we call behavioral modules: general inspection, general recognition and anxiety (see the results’ section for more details). Then, Linear Mixed Models (LMM) were used to assess the effects of sex, line, the interaction between the two and the fish ID as a random covariate on the scores each behavioral module, followed by Tukey post-hoc tests. These analyses were carried out in the statistical software Minitab ® version 17 (Minitab Inc., State Collage, PA, USA).

The remaining analyses were carried out in the statistical software R, version 4.0.4 [57]. To test if the behavioral modules are differently related for with each zebrafish line, we computed Pearson correlation matrices between the three PC scores across each line. All p-values were corrected for multiple testing with Benjamin and Hochberg’s method. Heatmaps were used for visual representation of the correlation matrices for each line. The packages “Hmisc” [58] and “ggplot2” [59] were used for computing the correlations and building the heatmaps, respectively. The quadratic assignment procedure (QAP) correlation test with 5000 permutations [60], was used to assess the association between any two correlation matrices between different zebrafish lines on UCINET 6 [61]. Given that the null hypothesis of the QAP test is that there is no association between matrices, a significant p-value indicates that the correlation matrices are similar.

To check whether the genetic distances between subjects (using their genetic data from the list of 132 SNPs) are structured by line or represent a uniform population, we performed a hierarchical clustering analysis, using the “philentropy” package [62]. We computed the jaccard distance between all subjects, which is the proportion of the similar genetic distances between subjects over the total genetic distances. With the genetic distances’ matrix, we performed the hierarchical clustering with complete-linkage, which calculates the maximum distance between clusters before merging. Then, we plotted the hierarchical cluster in a dendrogram using the “dendextend” package [63]. We found a structed population with 5 clusters, corresponding to 4 of the 6 different lines, with the 5^th^ cluster merging the TU and WIK lines together (see results section for more details). Therefore, we decided to include line as a covariate in the analyses of SNP-behavior associations (see below).

To assess the associations between genetic polymorphisms and behavior, we tested each of the 132 SNPs independently against each behavioral phenotype (the 7 behaviors and 3 PC scores). We did not include 3 zebrafish subjects in this analysis because their sample call rate was below 5%, meaning they lack genetic information for most SNPs. For the behaviors that followed a linear distribution (general inspection, general recognition, anxiety and edge-orienting) we used linear models (LM) implemented with the R “base” package. For the behaviors that were proportions (social tendency, social discrimination, social exploration, object discrimination, object exploration and thigmotaxis), we used generalized linear models (GLM) with beta regression implemented with the “betareg” package [64]. In all models, the behaviors were the response variables, SNP was the explanatory variable and line was a co-variate. SNPs were integers, where 1 represented the heterozygote case and 0 and 2 the homozygotes. For example, for SNP rs180151563, 0 represents the genotype AA, 1 the genotype CA and 2 the genotype CC. For some SNPs there were only two of the three possible conditions. Line represents the different origins of the zebrafish subjects that we tested. It was also an integer, varying between 1 and 6, where 1 represented the 5D line, 2 the AB line, 3 the LEO line, 4 the TL line, 5 the TU line and 6 the WIK line. For each statistical model, we used the summary() function in R to extract the p-value of the SNP, which was corrected for the line effect. Because we run 132 independent tests for each SNP, we corrected the p-values with the false discovery rate (FDR) adjustment method.

For some of the SNP-behavior associations that remained significant after FDR adjustment, we used the “ggplot2” [59] and “ggpubr” [65] packages to draw boxplots for the given phenotype, broken down by the SNP genotype. Over the boxplots, we added dot plots broken down by Line to help visualizing the Line effect on zebrafish behavior. For a more comprehensive comparison of the SNP-behavior associations, we plotted the significant associations by behavioral categories using Venn diagrams, with the “VennDiagram” package [66].

## Supporting information

Table S1 presents the results of one-sample t-tests that assess if scores of social tendency (i.e. preference for shoal over empty tank), as well as object and conspecific discrimination scores (i.e. preference between a novel and a familiar stimulus) were significantly different than chance for individuals of both sexes and for all lines tested (one-sample *t*-test: *μ ≠* 0.5, *P* < 0.05).

## Data Availability statement

Data has been deposited at the public repository Dryad (https://doi.org/10.5061/dryad.v15dv41wh).

## Acknowledgements

The authors thank Ibukun Akinrinade for extracting the DNA samples for the SNP analysis. This study was funded by Fundação para a Ciência e a Tecnologia (FCT, Portugal) through grants PTDC/BIA-ANM/0810/2014 and PTDC/BIA-COM/30627/2017 awarded to RFO and a PhD fellowship awarded to C.G. (SFRH/BD/132562/2017). All the genetic work was run at the Genomics Unit of Instituto Gulbenkian de Ciência which is funded by a FCT R&D unit grant (UIDB/04555/2020).

## Author Contributions

Conceptualization: CG, RFO

Data curation: CG, KK, SAMV

Formal analysis: KK, SAMV, TP, RFO

Funding acquisition: RFO, CG

Investigation: CG, MCT, KK, JC, RBL

Methodology: MCT, JC, RBL

Project administration: RFO

Supervision: RFO

Visualization: KK, SAMV, MCT, JC

Writing – original draft: CG, KK, RFO

Writing – review & editing: all authors

## References

1. Ward A, Webster M. Sociality: The Behaviour of Group-Living Animals. Springer-Verlag, NY. 2016.

2. Okuyama T. Social memory engram in the hippocampus. Neurosci Res. 2018; 129: 17–23.

3. Dölen G, Darvishzadeh A, Huang K, Malenka R. Social reward requires coordinated activity of nucleus accumbens oxytocin and serotonin. Nature. 2013; 501: 179–184.

4. Gunaydin LA, Grosenick L, Finkelstein JC, Kauvar IV, Fenno LE, Adhikari A, et al. Natural neural projection dynamics underlying social behavior. Cell. 2014; 157: 1535–1551.

5. Heyes C, Pearce JM. Not-so-social learning strategies. P R Soc B. 2015; 282: 20141709.

6. Varela SAM, Teles M, Oliveira RF. The correlated evolution of social competence and social cognition. Funct Ecol. 2020; 34(2): 332–343.

7. Bowen MT, Keats K, Kendig MD, Cakic V, Callaghan PD, McGregor IS. Aggregation in quads but not pairs of rats exposed to cat odor or bright light. Behav Processes. 2012; 90(3): 331–336.

8. Kleinhappel TK, Pike TW, Burman OHP. Stress-induced changes in group behaviour. Sci Rep. 2019; 9: 17200.

9. Bell A. Behavioural differences between individuals and two populations of stickleback (Gasterosteus aculeatus). J Evolution Biol. 2005; 18(2): 464–473.

10. Sören K, Frieder P, Maren B, TIlo K. The rewarding nature of social interactions. Front Behavl Neurosci. 2010; 4: 22.

11. Gunaydin L, Deisseroth K. Dopaminergic Dynamics Contributing to Social Behavior. Cold Spring Harb Sym. 2014; 79: 221–227.

12. Walsh JJ, Christoffel DJ, Heifets BD, Ben-Dor GA, Selimbeyoglu A, Hung LW, Deisseroth K, Malenka RC. 5-HT release in nucleus accumbens rescues social deficits in mouse autism model. Nature. 2018; 560(7720): 589–594.

13. Donaldson Z, Young L. Oxytocin, vasopressin, and the neurogenetics of sociality. Science. 2008; 322: 900–904.

14. Goodson J, Thompson R. Nonapeptide mechanisms of social cognition, behavior and species-specific social systems. Curr Opin Neurobiol. 2010; 20(6): 784–794.

15. Goodson J. Deconstructing sociality, social evolution and relevant nonapeptide functions. Psychoneuroendocrino. 2013; 38(4): 465–478.

16. Südhof T. Neuroligins and Neurexins Link Synaptic Function to Cognitive Disease. Nature. 2008; 455: 903–911.

17. Grayton M, Missler M, Collier D, Fernandes C. Altered Social Behaviours in Neurexin 1α Knockout Mice Resemble Core Symptoms in Neurodevelopmental Disorders. Plos One. 2013; 8(6): e67114.

18. Rabaneda L, Robles-Lanuza E, Nieto-González J, Scholl F. Neurexin Dysfunction in Adult Neurons Results in Autistic-like Behavior in Mice. Cell Reports. 2014; 8(2): 338–346.

19. Hörnberg H, Pérez-Garci E, Schreiner D, Hatstatt-Burklé L, Magara F, Baudouin S, et al. Rescue of oxytocin response and social behaviour in a mouse model of autism. Nature. 2020; 584: 252–256.

20. O’Connell LA, Hofmann HA. The Vertebrate Mesolimbic Reward System and Social Behavior Network: A Comparative Synthesis. J Comp Neurol. 2011; 519: 3599–3639.

21. O’Connell LA, Hofmann HA. Evolution of a vertebrate social decision-making network. Science. 2012; 336(6085): 1154–1157.

22. Tibbetts E, Dale J. Individual recognition: it is good to be different. Trends Ecol Evol. 2007; 22(10): 529–537.

23. Mrakovcic M, Haley L. Inbreeding depression in the Zebra fish *Brachydanio rerio*(Hamilton Buchanan). J Fish Biol. 1979; 15(3): 323–327.

24. Nasiadka A, Clark M. Zebrafish breeding in the laboratory environment. ILAR J. 2012; 53(2): 161–168.

25. Brown KH, Dobrinski KP, Lee AS, Gokcumen O, Mills RE, Shi X, et al. Extensive genetic diversity and substructuring among zebrafish strains revealed through copy number variant analysis. P Natl Acad Sci USA. 2012; 109(2): 529–534.

26. Balik-Meisner M, Truong L, Scholl EH, Tanguay RL, Reif DM. Population genetic diversity in zebrafish lines. Mamm Genome. 2018; 29: 90–100.

27. Oswald M, Robison BD. Strain-specific alteration of zebrafish feeding behavior in response to aversive stimuli. Can J Zool. 2008; 86(10): 1085–1094.

28. Egan RJ, Bergner CL, Hart PC, Cachat JM, Canavello PR, Elegante MF, et al. Understanding behavioral and physiological phenotypes of stress and anxiety in zebrafish. Behav Brain Res. 2009; 205(1): 38–44.

29. Sackerman J, Donegan JJ, Cunningham CS, Nguyen NN, Lawless K, Long A, Benno RH, Gould GG. Zebrafish Behavior in Novel Environments: Effects of Acute Exposure to Anxiolytic Compounds and Choice of *Danio rerio* Line. J Comp Psychol. 2010; 23(1): 43–61.

30. Barba-Escobedo P, Gould G. Visual social preferences of lone zebrafish in a novel environment: strain and anxiolytic effects. Genes, Brain and Behav. 2012; 11(3): 366–373.

31. Lange M, Neuzeret F, Fabreges B, Froc C, Bedu S, Bally-Cuif L, Norton WHJ. Inter-Individual and Inter-Strain Variations in Zebrafish Locomotor Ontogeny. PLoS One. 2013; 8(8): e70172.

32. Mahabir S, Chatterjee D, Buske C, Gerlai R. Maturation of shoaling in two zebrafish strains: A behavioral and neurochemical analysis. Behav Brain Res. 2013; 247: 1–8.

33. Maximino C, Puty B, Benzecry R, Araújo J, Lima MG, Batista E de JO, et al.. Role of serotonin in zebrafish (*Danio rerio*) anxiety: Relationship with serotonin levels and effect of buspirone, WAY 100635, SB 224289, fluoxetine and para-chlorophenylalanine (pCPA) in two behavioral models. Neuropharmacology. 2013; 71: 83–97.

34. Vignet C, Bégout M-L, Péan S, Lyphout L, Leguay D, Cousin X. Systematic Screening of Behavioral Responses in Two Zebrafish Strains. Zebrafish. 2013; 10(3): 365–375.

35. Vital C, Martins EP. Strain differences in zebrafish (*Danio rerio*) social roles and their impact on group task performance. J Comp Psychol. 2011; 125(3): 278–285.

36. Liu X, Guo N, Lin J, Zhang Y, Chen XQ, Li S, He L, Li Q. Strain-dependent differential behavioral responses of zebrafish larvae to acute MK-801 treatment. Pharmacol Biochem Behav. 2014; 127: 82–9.

37. Gorissen M, Manuel R, Pelgrim TNM, Mes W, de Wolf MJS, Zethof J, Flik G, van den Bos R. Differences in inhibitory avoidance, cortisol and brain gene expression in TL and AB zebrafish. Genes Brain Behav. 2015; 14(5): 428–438.

38. Séguret A, Collignon B, Halloy J. Strain differences in the collective behaviour of zebrafish (*Danio rerio*) in heterogeneous environment. Roy Soc Open Sci. 2016; 3(10): 160451.

39. Ribeiro D, Nunes AR, Gliksberg M, Anbalagan S, Levkowitz G, Oliveira RF. Oxytocin receptor signalling modulates novelty recognition but not social preference in zebrafish. J Neuroendocrinol. 2020; 32: e12834.

40. Groenewoud F, Frommen J, Josi D, Tanaka H, Jungwirth A, Taborsky M. Predation risk drives social complexity in cooperative breeders. P Natl Acad Sci USA. 2016; 113(15): 4104–4109.

41. Ferguson JN, Aldag JM, Insel TR, Young LJ. Oxytocin in the medial amygdala is essential for social recognition in the mouse. J Neurosci. 2001; 21(20): 8278–85.

42. Resendez SL, Namboodiri VMK, Otis JM, Eckman LEH, Rodriguez-Romaguera J, Ung RL, et al. Social Stimuli Induce Activation of Oxytocin Neurons Within the Paraventricular Nucleus of the Hypothalamus to Promote Social Behavior in Male Mice. J Neurosci. 2020; 40(11): 2282–2295.

43. Dochtermann N, Dingemanse N. Behavioral syndromes as evolutionary constraints. Behav Ecol. 2013; 24(4): 806–811.

44. Saltz J, Hessel F, Kelly M. Trait correlations in the genomics era. Trends Ecol Evol. 2017; 32(4): 279–290.

45. Royauté R, Hedrick A, Dochtermann NA. Behavioural syndromes shape evolutionary trajectories via conserved genetic architecture. P R Soc B. 2020; 287: 20200183.

46. Wright D, Nakamichi R, Krause J, Butlin RK. QTL Analysis of Behavioral and Morphological Differentiation Between Wild and Laboratory Zebrafish (*Danio rerio*). Behav Genet. 2006; 36: 271.

47. Streisinger G, Walker C, Dower N, Knauber D, Singer F. Production of clones of homozygous diploid zebra fish (*Brachydanio rerio*). Nature. 1981; 291: 293–296.

48. Chakrabarti S, Streisinger G, Singer F, Walker C. Frequency of gamma-ray induced specific locus and recessive lethal mutations in mature germ cells of the zebrafish, *Brachydanio rerio*. Genetics. 1983; 103(1): 109–123.

49. Mullins MC, Hammerschmidt M, Haffter P, Nusslein-Volhard C. Large-scale mutagenesis in the zebrafish: in search of genes controlling development in a vertebrate. Curr Biol. 1994; 4(3): 189–202.

50. Nusslein-Volhard C. In: Zebrafish: A Practical Approach. Dahm R, editor. New York: Oxford Univeristy Press. 2002.

51. Rauch G-J, Granato M, Haffter P. A polymorphic zebrafish line for genetic mapping using SSLPs on high-percentage agarose gels. Technical Tips Online. 1997; 2: 148–150.

52. Kent ML, Buchner C, Watral VG, Sanders JL, LaDu J, Peterson TS, Tanguay RL. Development and maintenance of a specific pathogen-free (SPF) zebrafish research facility for Pseudoloma neurophilia. Dis Aquat Organ. 2011; 95(1): 73–9.

53. Frankel J. Inheritance of spotting in the leopard Danio. J Hered. 1979; 70(4): 287–288.

54. Haffter P, Odenthal J, Mullis M., Lin S, Farrell MJ, Vogelsang E, et al. Mutations affecting pigmentation and shape of the adult zebrafish. Dev Genes Evol. 1996; 206: 260–276.

55. Watanabe M, Iwashita M, Ishii M, Kurachi Y, Kawakami A, Kondo S, Okada N. Spot pattern of leopard Danio is caused by mutation in the zebrafish connexin41.8 gene. EMBO Reports. 2006; 7(9): 893–897.

56. Kalueff AV, Gebhardt M, Stewart AM, Cachat JM, Brimmer M, Chawla JS, et al. and the Zebrafish Neuroscience Research Consortium (ZNRC). Towards a comprehensive catalog of zebrafish behavior 1.0 and beyond. Zebrafish. 2013; 10(1): 70–86.

57. R Core Team. R: A language and environment for statistical computing. R Foundation for Statistical Computing, Vienna, Austria. URL. 2021. https://www.R-project.org/.

58. Harrell F E Jr, with contributions from Charles Dupont and many others. (2020). Hmisc: Harrell Miscellaneous. R package version 4.4-0. 2020. https://CRAN.R-project.org/package=Hmisc.

59. Wickham H. ggplot2: Elegant Graphics for Data Analysis. Springer-Verlag New York. 2016.

60. Borgatti SP, Everett MG, Johnson JC. Analyzing social networks. Beverley Hills, CA: Sage Publications. 2013.

61. Borgatti SP, Everett MG, Freeman LC. Ucinet for Windows: Software for Social Network Analysis. Harvard, MA: Analytic Technologies. 2002.

62. Drost H-G. Philentropy: Information Theory and Distance Quantification with R. Journal of Open Source Software. 2018; 3(26): 765.

63. Galili T. Dendextend: an R package for visualizing, adjusting, and comparing trees of hierarchical clustering. Bioinformatics. 2015; 31(22): 3718–3720.

64. Cribari-Neto F, Zeileis A. Beta Regression in R. J Stat Softw. 2010; 34(2): 1–24.

65. Kassambara A. ggpubr: ‘ggplot2’ Based Publication Ready Plots. R package version 0.4.0. 2020. https://CRAN.R-project.org/package=ggpubr.

66. Chen H. Venn Diagram: Generate High-Resolution Venn and Euler Plots. R package version 1.6.20. 2018. https://CRAN.R-project.org/package=VennDiagram

